# Harnessing Genome Representation Learning for Decoding Phage-Host Interactions

**DOI:** 10.1101/2024.03.12.584599

**Authors:** Sumanth Badam, Shrisha Rao

## Abstract

Accurate prediction of the phages that target a bacterial host plays an important role in combating anti-microbial resistance. Our work explores the power of deep neural networks, convolutional neural networks, and pre-trained large DNA/protein language models to predict the host for a given phage. This work mainly uses the data provided by Gonzales et al. that contains receptor-binding protein sequences of the phages and the target host genus. We used pre-trained language models to obtain the dense representations of protein/nucleotide sequences to train a deep neural network to predict the target host genus. Additionally, convolutional neural networks were trained on one-hot encoding of nucleotide sequences to predict the target host genus. We achieved a weighted F1-score of 73.76% outperforming state-of-the-art models with an improvement of around 11% by using the protein language model ESM-1b.

The data and the source code are available at https://github.com/sumanth2002629/Bacteriophage-Research.

## Introduction

Bacterial infections, an age-old menace to human society, were considered to have been brought under control thanks to antibiotics such as penicillin. However dangerous bacteria have evolved in a way to lessen their susceptibility, and as a result, many antibiotics are no longer as potent as they once were. Additionally, creating and implementing new antibiotic treatments remains a laborious and challenging process that involves identifying appropriate chemical compounds, obtaining regulatory clearances, and setting up production systems. In light of this, bacteriophages, which are viruses that target bacteria, are being thoroughly studied as potential substitutes for traditional approaches to treating bacterial infections. This work is a significant improvement in phage-host interaction prediction harnessing the power of large DNA/Protein language models and deep neural networks. We have made use of the dataset that is made available by Gonzales et al. (Gonzales et al., 2023). The dataset consists of many attributes of phages including the receptor-binding protein (RBP) sequence and the nucleotide sequence of phages with the target variable as the host genus. Gonzales et al. (Gonzales et al., 2023) have collected the data from GenBank via INPHARED (Cook et al., 2021) and trained an XGBoost model on the embeddings of receptor binding protein (RBP) sequences obtained from various protein language models. The weighted F1-score that they obtained in predicting the host given an RBP sequence of a phage is 62.95%. The shortcoming of their work is that they haven’t made use of the available nucleotide sequences for this task. They have also not experimented with deep learning methods which are capable of recognizing complex patterns.

We overcome the shortcomings of the previous work by making use of the phage’s receptor-binding nucleotide sequences in two ways. Firstly, we have encoded the nucleotide sequence of the phage in a onehot fashion. Then we trained a 1-dimensional convolutional neural network (that has one convolutional layer followed by a fully connected layer) on the onehot encoded data to predict the host genus. The dimension of the input layer is four times the length of the longest nucleotide sequence which is 19044 (4*×*4761). This model performs with a weighted F1score of **72.73**% which is in an improvement of **10**% in the weighted F1-score compared to the earlier work (Gonzales et al., 2023). Table 7 shows the results in more detail.

The other way in which we make use of the nucleotide (DNA) sequences is by obtaining the embeddings from a DNA language model called DNA-BERT2 (Zhou et al., 2023) which is a foundational model trained on a large-scale multi-species genome. It improves the efficiency and effectiveness of DNABERT (Ji et al., 2021) by incorporating techniques like replacing *k*mer tokenization with BPE, and positional embedding with Attention with Linear Bias (ALiBi). By training a neural network on the embeddings from DNA-BERT2 (Zhou et al., 2023), we obtain a competitive performance of **60.23**% in terms of weighted F1 score. Table 6 shows the results in more detail.

We have made use of the protein embeddings provided by Gonzales et al. (Gonzales et al., 2023) by training a more complex model to predict the host genus. We have trained a fully connected 3-layer neural network on the protein embeddings obtained from various protein language models. The input layer dimension of the neural networks varies according to the length of the embedding from a protein language model. A neural network that is trained on the embeddings from ESM-1b (Rives et al., 2021) performs with a weighted F1 score of 73.76%, which is an improvement of about 11% in the weighted F1-score compared to the earlier work (Gonzales et al., 2023). Table 5 shows the results in more detail. We have also tried concatenating the protein and the DNA embeddings which are passed as inputs to the neural network. However, this does not give a better performance compared to the protein embeddings alone.

Our novel contributions are summarised below:

- Fully connected neural network trained on the protein embeddings obtained from various protein language models surpasses the performance of state-of-the-art by around 11% giving a weighted F1-score of 73.76%
- Fully connected neural network trained on the DNA embeddings obtained from DNA-BERT2 gives a competitive performance of 60.23% in terms of weighted F1 score.
- Convolutional neural network trained on the one-hot encodings obtained from the nucleotide (DNA) sequences gives an improvement over the state-of-the-art by around 10% (72.73%).

The structure of the rest of this article is as follows: Section- presents the background of various works related to phage-host interaction prediction. A detailed evolution of the studies ranging from in vitro experiments to the use of transformer-based models is described in detail. Section- explains the data preparation, model architectures, and the evaluation methodology in detail. Section-D contains the four main result objects that correspond to the four experiments done. Lastly, in Section-D, we conclude by providing various future directions for this work.

## Background

In vitro trials to choose the potential phages that target a host are expensive and time-consuming. The improvement in the high throughput sequencing technologies resulted in increased omic data. This helped researchers to make use of the computational methods for phage host interaction prediction. These are mainly categorized into alignment-based methods (Altschul et al., 1990; Zielezinski et al., 2021) and alignment-free methods (Villarroel et al., 2016; Galiez et al., 2017). Alignment-based methods study the similarity between the omic sequences of the phages and the host whereas alignment-free methods use sequence composition features such as codon usage bias and oligonucleotide frequency.

Machine Learning algorithms were also explored to predict the phage-host interaction. Most of them consider the entire proteome sequences as the feature sets for both binary and multiclass classification (Leite et al., 2018; Zhou et al., 2022; Li et al., 2020; Li and Zhang, 2022; Tan et al., 2022; Ruohan et al., 2022) when only specific proteins called receptor binding proteins are involved in the phage-host interaction. These proteins are present at the end of the phage’s tail which absorbs to the host surface (Nobrega et al., 2018). Our work makes use of the receptor-binding proteins as feature vectors with the target class as the host genus. Recent applications of representation learning, which turns unprocessed biological sequences into dense vectors in high-dimensional space, include the prediction of protein function (Nobrega et al., 2018), the identification of succinylation sites (Pokharel et al., 2022), and the conservation of sequences (Marquet et al., 2022). Transformer-based large protein-language models like ESM (Rives et al., 2021) and ProtTrans (Elnaggar et al., 2021) have been used to capture the features that are relevant to phage-host specificity. The evolution of studying the phage-host interaction prediction depended on the state-of-the-art technology of that time. In machine learning, transformer-based architectures are the current state-of-the-art and we utilize the embeddings from pre-trained transformer-based models for our study.

In the recent work done by Gonzales et al. (Gonzales et al., 2023), they use various protein language models to obtain the embeddings from receptor-binding protein sequences and they use those embeddings to train an XGBoost model that predicts the target host. In our work, we only focus on the receptor-binding protein/nucleotide sequences which determine the phagehost interaction. We made use of representation learning by transforming the nucleotide sequences into a higher dimensional space using DNABERT2 (Zhou et al., 2023). We have used the embeddings of protein sequences provided by Gonzales et al. (Gonzales et al., 2023). As per our knowledge, we are the first to make use of convolutional neural networks on receptor binding protein (RBP) sequences of phages to study phage-host interaction.

## Data and Methods

We extensively made use of data that is made available by Gonzales et al. (Gonzales et al., 2023). where they have used INPHARED (Cook et al., 2021) to extract phage sequences from GenBank (Benson et al., 2005) and performed necessary pre-processing. It consists of various attributes of phages mainly receptor-binding nucleotide sequences and receptor-binding protein sequences. It also contains the protein embeddings obtained upon passing RBP sequences as an input to the protein language model. Various pre-trained protein language models like ESM (Rives et al., 2021), ESM-1b (Rives et al., 2021), ProtBert (Elnaggar et al., 2021), ProtXLNet (Elnaggar et al., 2021), ProtAlbert (Elnaggar et al., 2021), ProtT5 (Elnaggar et al., 2021), and SeqVec (Heinzinger et al., 2019) were used to obtain RBP embeddings. The target variable Host contains the host genus.

We used DNA-BERT2 (Zhou et al., 2023) a DNA language model, to obtain DNA embeddings of receptor-binding nucleotide sequences to study the efficacy of DNA embeddings in phage-host interaction prediction.

To study the efficacy of Convolutional Neural Networks (CNNs)(LeCun et al., 2015) in phage host interaction prediction, we trained the network with the one-hot encoding of the DNA sequences. Onehot encoding was obtained by encoding A as 1000, C as 0100, G as 0010, and T as 0001. For example, one-hot encoding of the sequence ATTTACGG is 10000001000100011000010000100010. The length of the input vector after encoding would be four times the length of the longest nucleotide sequence. The entries are padded with 0’s for the shorter sequences. The advantage of one-hot encoding is that it allows the model to learn more easily and effectively. This is because each input value is represented as a binary vector, where only a single element of the vector is set to 1 and the rest are set to 0. This makes it easier for the model to learn the relationship between the input values and the target output because the model can easily distinguish between the different input values based on their unique binary representation.

### A. Train-Test split

For each of the experiments, we have the corresponding feature vector and the target variable Host. In total, we have 24,752 entries with 232 hosts. Out of 232 hosts, the top 58 hosts that are more frequent in the dataset are associated with 96.02% of the dataset entries. Let *D*_1_ be the dataset entries corresponding to the top 58 (25%) of the hosts and *D*_2_ be the dataset with the other 75% of the hosts.

*D*_1_ is partitioned into train and test datasets such that the training dataset contains 70% of *D*_1_ and the test dataset contains 30% of *D*_1_. The split was performed in such a way that the train and test datasets have the same proportion (70%-30%) for every host. Then we append *D*_2_ to the test dataset by marking all the hosts of *D*_2_ as *others*.

Now the test and train datasets have 16,636 and 8,116 entries respectively. The hosts were encoded into integers 0 to 57 for the top 58 hosts and the *others* class was encoded as -1.

### B. ANNs to train on embeddings from Protein/DNA language models

We trained a fully connected neural network to predict the target host. Our model has 4 layers with one input layer, two hidden layers, and an output layer. There is a ReLU activation layer after each of the hidden layers. The loss function used is binary crossentropy loss.

We use two different neural networks, one for the protein embeddings and one for the DNA embeddings. Both the networks have same number of nodes in hidden layers and the output layer. However, the number of nodes in the input layer depends on the embedding size. In the data provided by Gonzales et al. (Gonzales et al., 2023), the size of protein embeddings is 1024 (for any protein language model). Whereas, DNA-BERT2 outputs an embedding of size 768. The model architectures are clearly described in Table 1 and Table 2.

**Table 1.** Architecture of fully connected neural network that takes protein embedding as an input.

| Layers | Type | Shape of Output | Parameters |
| --- | --- | --- | --- |
| dense_1 | Dense | (None,512) | 393728 |
| relu_1 | Activation | (None,512) | 0 |
| dense_2 | Dense | (None,256) | 262656 |
| relu_2 | Activation | (None,256) | 0 |
| dense_3 | Dense | (None,58) | 29754 |

**Table 2.** Architecture of fully connected neural network that takes DNA embedding as an input.

| Layers | Type | Shape of Output | Parameters |
| --- | --- | --- | --- |
| dense_1 | Dense | (None,512) | 524800 |
| relu_1 | Activation | (None,512) | 0 |
| dense_2 | Dense | (None,256) | 262656 |
| relu_2 | Activation | (None,256) | 0 |
| dense_3 | Dense | (None,58) | 29754 |

### C. CNNs to train on the one-hot encoding of DNA sequences

We trained a convolutional neural network (CNN) to predict the target host. The CNN, which was trained on the one-hot encoding of the DNA sequence, has one 1D convolutional layer with one filter, followed by a fully connected layer. The loss function used is binary cross-entropy loss. The CNN architecture is clearly described in Table 3.

**Table 3.** Architecture of convolutional neural network.

| Layers | Filters | Stride | Filter Size | Output shape |
| --- | --- | --- | --- | --- |
| conv_1 | 1 | 1 | $6 \times 1$ | (None,19039) |
| relu_1 | 0 | - | - | (None,19039) |
| dense_1 | - | - | - | (None, 58) |

### D. Evaluation Methodology

We have a threshold parameter *k* which determines the confidence of the prediction. Let *p*_1_ and *p*_2_ be the first and second highest probabilities of the posterior probability vector. If *p*_1_*−p*_2_ *≥ k*, we classify the input as a class corresponding to *p*_1_. Otherwise, the input is classified as *others*(−1).

**Figure 1.**
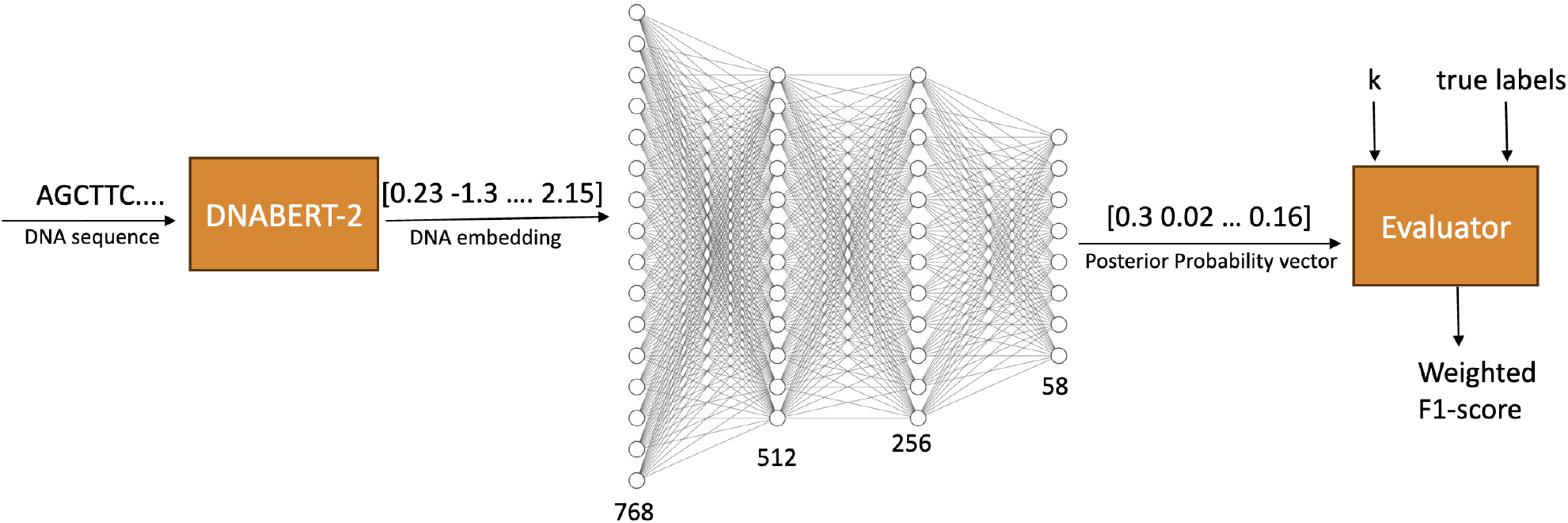
Methodology of our study: DNABERT-2 generates the embeddings by taking DNA sequence as an input which are then passed to a fully connected neural network to output the posterior probability vector. True labels along with the posterior probability vector and the threshold k are passed to the evaluator to compute the F1 score.

Once, we have the predicted output vector and the true labels, the weighted F1 score is calculated as follows:

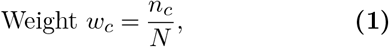

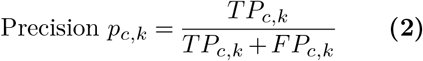

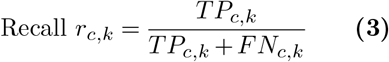

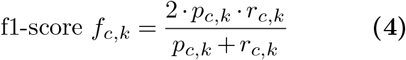

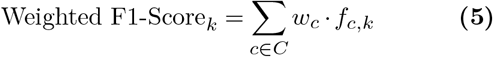

Formally, *N* is the total number of test samples, *n*_*c*_ is the total number of test samples of class *c ∈ C* where *C* is the set of all classes including *others*. Weighted F1-Score for a given threshold *k* is calculated as shown in 5.

## Results

We evaluated our models for different values of threshold (*k*) ranging from 0.6 to 1.0 in steps of 0.1. We have four main result objects by evaluating CNN and two fully connected networks which were trained on protein and DNA sequences.

We compared our results with the state-of-the-art performance in the work done by Gonzales et al. (Gonzales et al., 2023). We got a weighted F1 score of 73.76% on training a fully connected network with the embeddings from ESM-1b which is around 11% greater than the f1-score that was obtained by Gonzales et al. (Gonzales et al., 2023). Our results in Table 5 for different protein language models can be compared with the results of the work done by Gonzales et al. (Gonzales et al., 2023) in Table 4.

**Table 4.**
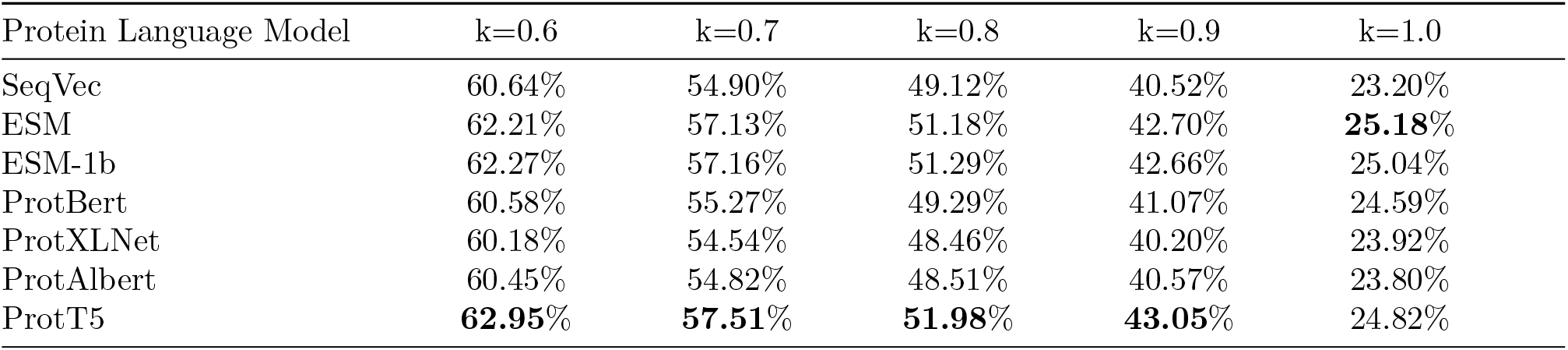
Performance of the XGBoost model trained on protein embeddings in terms of weighted F1 score in the work done by Gonzales et al. (Gonzales et al., 2023). Each row corresponds to the performance of the XGBoost model on the embeddings from a protein language model for various thresholds (k). (The highest scores are in bold)

**Table 5.**
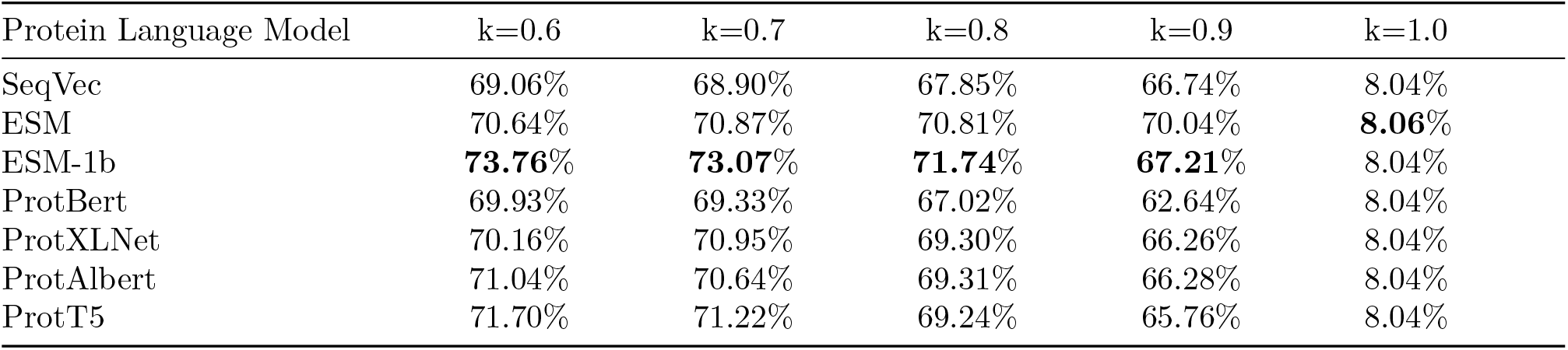
Performance of the fully connected neural network trained on protein embeddings in terms of weighted F1 score in this work. Each row corresponds to the performance of the fully connected neural network on the embeddings from a protein language model for various thresholds(k). (The highest scores are in bold).

Table 6 shows that training a fully connected network on DNA embeddings from DNABERT2 gives a competitive performance of 60.82% (weighted f1score) with respect to the state-of-the-art.

**Table 6.**
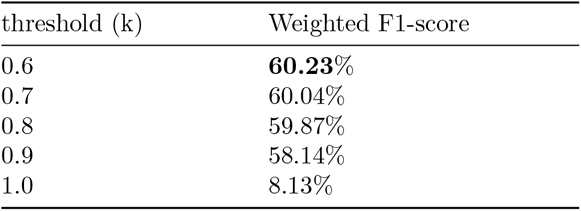
Performance of a fully connected neural network that was trained on the embeddings from DNABERT2.(The highest score is in bold)

| threshold (k) | Weighted F1-score |
| --- | --- |
| 0.6 | **60.23%** |
| 0.7 | 60.04% |
| 0.8 | 59.87% |
| 0.9 | 58.14% |
| 1.0 | 8.13% |

Table 7 shows that training a convolutional neural network on the one-hot encoding of the DNA sequences gives a performance of 72.73% (weighted f1score) which is around 10% more than the f1-score that was obtained by Gonzales et al. (Gonzales et al., 2023).

**Table 7.** Performance of a convolutional neural network that was trained on the one-hot encoding of the DNA sequences. (The highest score is in bold)

| threshold(k) | Weighted F1-score |
| --- | --- |
| 0.6 | **72.73%** |
| 0.7 | 71.66% |
| 0.8 | 70.50% |
| 0.9 | 67.71% |
| 1.0 | 8.03% |

We have also trained a neural network by combining the embeddings of ProtT5 and ESM-1b (whose embeddings gave the best results in Table 5). The highest weighted F1-score is 71.70% for k=0.6 which is around 9% more than the previous work. Table 8 shows the results in detail.

**Table 8.** Performance of a fully connected neural network that was trained on the combined embeddings of ProtT5 and ESM-1b.(The highest score is in bold)

| threshold (k) | Weighted F1-score |
| --- | --- |
| 0.6 | **71.70%** |
| 0.7 | 71.22% |
| 0.8 | 69.24% |
| 0.9 | 65.76% |
| 1.0 | 8.04% |

## Conclusion

In this study, we have used the dataset that maps the RBP sequences to the host genus and performed several experiments that have resulted in improvement of the performance in predicting the host. Many other experiments that we have planned can be performed with extra computational resources.

Future scope by utilizing more computational resources include:

- Fine-tuning DNABERT-2 on the DNA sequences of phages to obtain better embeddings. As DNABERT-2 is a transformer-based model, it requires more resources to fine-tune the model.
- Experimenting with various architectures of CNNs with increasing the number of layers. It also requires more resources as the input size of one-hot encoding of DNA sequence is around twenty thousand.
- One can also experiment by encoding the protein sequences in a one-hot fashion. As there are 20 amino acids, the encoding of a protein sequence will become 20 times the length of the raw protein sequence.

Further aspects of the future scope include investigating other formulations of phage-host interaction prediction. For example, it can formulated as a binary classification problem in which given the phage and host information, the model predicts if the phage can target the host or not. Or given the genome sequences of the host, one can predict which phage is most likely to target the host and eventually destroy it. As CNN performed better, experimenting with RNN/LSTM-based architectures which are usually used to deal with sequence data is also an option. Identification of bacterial hosts at the strain level could be done by conducting some lab work or using the dataset that has the host strain as the target variable which is out of the scope of this work.

To provide more reasoning on why the CNN model has a high performance, we can look at DeepLIFT (Shrikumar et al., 2017) attribution scores for the first CNN layer. DeepLIFT attribution plot highlights the particular sub-sequence to be important which can be used to study the similarity between the highlighted sequences of phages that target the same host. One can also look into studying the interpretability of DNA embeddings.

## Notes

https://github.com/sumanth2002629/Bacteriophage-Research

### Competing Interest Statement

The authors have declared no competing interest.

